# Polyphenol oxidase depletion in *Nicotiana benthamiana* enhances recombinant protein purification and preserves native protein integrity

**DOI:** 10.1101/2025.09.28.679031

**Authors:** Kaijie Zheng, Farnusch Kaschani, Emma C. Watts, Markus Kaiser, Renier A. L. van der Hoorn

## Abstract

Agroinfiltration of *Nicotiana benthamiana* is widely used for recombinant protein production in plant science and molecular pharming, but enzymatic browning and native protein crosslinking during extraction may limit protein integrity and purification efficiency. We generated genome-edited *N. benthamiana* lines lacking two polyphenol oxidases (PPOs) and analysed protein integrity, enzymatic activity profiles, and recombinant protein purification under non-denaturing extraction conditions. PPO-deficient plants showed reduced browning and native protein crosslinking, preserved endogenous proteins at their predicted molecular weights, displayed increased detectable enzyme activities, and achieved a significantly higher recovery and improved purity of a transiently expressed recombinant protein. These findings identify PPO-mediated oxidation as a major bottleneck during protein extraction and demonstrate that PPO depletion enhances recombinant protein purification while preserving native protein integrity.

## Introduction

Agroinfiltration of *Nicotiana benthamiana* has become one of the most widely used transient expression platforms in plant science and molecular pharming, owing to its speed, scalability, low production cost, and ability to produce complex eukaryotic proteins with post-translational modifications (Beritza et al., 2024a; Lawson et al., 2025; Bally et al., 2020; Golubova et al., 2024). This system is routinely used to study protein–protein interactions (Win et al., 2011), enzymatic activities (Dudley et al., 2022), protein structure (Lawson et al., 2025) and subcellular localization (Alamos et al., 2025), and has been successfully deployed for the production of vaccines (Ward et al., 2020; Ward et al., 2021), antibodies (Shanmugaraj et al., 2020; de Taeye SW et al., 2025), and other biopharmaceuticals (Busold et al., 2022; Kaldis et al., 2023; VanderBurgt et al., 2023; Busold et al., 2024), including plant-based COVID-19 vaccines (Charland et al., 2022; Hager et al., 2022).

Despite these advantages, the purification of recombinant proteins from agroinfiltrated *N. benthamiana* leaves remains a major bottleneck (Buyel et al., 2025). Leaf homogenization inevitably disrupts cellular compartmentalization, releasing phenolic compounds, chlorophyll, and a diverse array of endogenous enzymes that can compromise protein integrity, activity, and purity (Jutras et al., 2020; Daduang et al., 2025; Buyel et al., 2025; Lawson et al., 2025). One of the most prominent and widely observed problems during protein extraction from plant tissues is enzymatic browning, a process driven primarily by polyphenol oxidases (PPOs). PPOs are copper-dependent oxidoreductases localized to chloroplasts that catalyze the oxidation of monophenols and o-diphenols into highly reactive o-quinones (Boeckx et al., 2015). These quinones can polymerize into brown, melanin-like pigments and react nonspecifically with nucleophilic residues on proteins, leading to covalent crosslinking and loss of solubility (Sommer et al., 1994; Sui et al., 2023; Tang et al., 2023).

In fruits and vegetables, PPO-mediated browning is well known to reduce food quality, and extensive efforts have been devoted to its suppression through breeding, genome editing, and chemical inhibition (Terefe et al., 2014; González et al., 2020; Ren et al., 2024; Zou et al., 2025). In contrast, the impact of PPO activity on recombinant protein extraction from plant expression systems has received comparatively little attention. Our recent work using virus-induced gene silencing (VIGS) demonstrated that transient depletion of PPO transcripts in *N. benthamiana* suppresses enzymatic browning and native protein crosslinking in leaf extracts, resulting in greener extracts and improved protein recovery (Mahadevan et al., 2025). Notably, PPO depletion prevented the formation of high-molecular-weight complexes of the large subunit of ribulose-1,5-bisphosphate carboxylase/oxygenase (RbcL), suggesting that PPO activity drives extensive native protein crosslinking during extraction (Mahadevan et al., 2025).

These observations raise important questions regarding the broader consequences of PPO activity for recombinant protein production and biochemical analyses. First, it remains unclear whether stable genetic removal of PPO activity confers similar or greater benefits compared to transient silencing approaches. Second, the extent to which PPO-mediated crosslinking affects endogenous enzymes and their measurable activities has not been systematically investigated. Third, it is unknown whether PPO depletion improves not only protein yield but also the purity of recombinant proteins purified under non-denaturing conditions, which is essential for functional, structural, and interaction studies.

Here, we address these questions by generating and characterizing genome-edited *N. benthamiana* lines lacking the two major *PPO* genes. We show that these *ppo* double knockout lines grow normally, exhibit reduced enzymatic browning and native protein crosslinking, preserve endogenous proteins at their predicted molecular weights, and display increased detectable enzymatic activities. Importantly, we demonstrate that PPO depletion substantially improves the yield and purity of a transiently expressed recombinant protein purified from total leaf extracts. Together, our results establish PPO knockout plants as a powerful resource for both plant science and molecular pharming applications.

## Materials and Methods

### Plants materials and growth conditions

*N. benthamiana* LAB was transformed with a T-DNA carrying Cas9 and two single guide RNAs targeting each gene (**Supplemental Table S1**). Two independent CRISPR mutants were identified as homozygous by PCR and sequencing using sequencing primers listed in **Supplemental Table S1**. The plants were maintained in a controlled growth room environment at 22 °C. The plants were exposed to a 16-h light/8-h dark photoperiod, under an illumination intensity of 2000 cd·sr·m⁻².

### Agroinfiltration

*Agrobacterium tumefaciens* GV3101 (pMP90) was used for agroinfiltration. Agrobacterium cultures were grown overnight in LB medium supplemented with 50 μg/ml kanamycin, 10 μg/ml gentamicin and 50 µg/ml rifampicin at 28 °C with shaking at 200 rpm. For transient protein expression assays, overnight cultures of Agrobacterium strains carrying binary vectors listed in **Supplemental Table S2** were harvested by centrifugation. Bacterial pellets were resuspended in infiltration buffer (10 mM MgCl₂, 10 mM MES, pH 5.6, 150 μM acetosyringone) and mixed at a 1:1 ratio with Agrobacterium harboring the silencing suppressor p19, with each strain adjusted to an OD₆₀₀ of 0.5. After incubation at 28 °C for 1 h, bacterial suspensions were infiltrated into leaves of 4-week-old *Nicotiana benthamiana* plants. Inoculated plants were maintained in a growth chamber until further use.

### Molecular cloning

To generate a plasmid encoding PPO1 flanked by FLAG and His tags, total RNA was extracted from leaves of ∼4-week-old wild-type *Nicotiana benthamiana* plants (FastPure Universal Plant Total RNA Isolation Kit, RC411-01, Vazyme), and the full-length open reading frame (ORF) of *PPO1* was amplified (Q5® Hot Start High-Fidelity DNA Polymerase, M0493S, NEB) by reverse transcription PCR (RT-PCR). The amplified ORF was subsequently recombined into a vector containing the 35S promoter, an N-terminal FLAG tag (Zheng et al., 2024), and a C-terminal His tag using ClonExpress Ultra One Step Cloning Kit V2 (C116-02, Vazyme), resulting in pKZ205.

### SDS-PAGE and Western blot

For total protein extraction from *Nicotiana benthamiana* leaves, samples were rapidly frozen in liquid nitrogen and ground to a fine powder with glass beads using a homogenizer. Depending on the experimental requirements, TBS, PBS or 2× gel loading buffer was added. When total protein was extracted using TBS or PBS, 4× gel loading buffer was subsequently added for SDS–PAGE analysis. After addition of the loading buffer, samples were denatured at 98 °C for 5 min and then loaded onto 12% SDS–PAGE gels and electrophoresed at 180 V in Invitrogen Novex vertical gel tanks. PageRuler™ Prestained Protein Ladder, 10 to 180 kDa (26616, Thermo Scientific™) was used as the protein ladder. Proteins were transferred to PVDF membranes using a BIO-RAD transfer apparatus and kit (Trans-Blot Turbo RTA Midi 0.45 µm LF PVDF Transfer Kit, 1704275), following the manufacturer’s instructions. Membranes were blocked in 5% skim milk in PBS-T (PBS tablets, 524650, Merck; 0.1% Tween-20, P1379, Merck) at room temperature for 1 h and then incubated overnight at 4 °C with primary antibodies against PPO (MBS9458497, MyBioSource), ALD (AS08294, Agrisera) and SHMT (AS05075, Agrisera) and GFP (Ab6663, Abcam), diluted 1:5000 in PBS-T. Primary antibody against RbcL (AS03037, Agrisera) was diluted 1:10000 in PBS-T. For PPO detection, membranes were further incubated with goat anti-mouse secondary antibody. Membranes were washed three times with PBS-T for 5 min each, followed by one wash with PBS for 5 min. Signals were developed using SuperSignal™ West Femto Maximum Sensitivity Substrate (34096, Thermo Scientific™) and detected with ImageQuant® LAS-4000 imager (GE Healthcare, Healthcare Life Sciences, Little Chalfont, UK).

### Serine hydrolase activity profiling

Total proteins were extracted from WT and *ppo* mutant plants using 50 mM sodium acetate (NaAc) buffer (pH 6.0), with minor modifications to the previously described procedure (Kaschani et al., 2009). Briefly, 25 μl of total protein extract was incubated with 0.2 μM FP-TAMRA (Thermo Scientific™) and adjusted to a final volume of 500 μl with 50 mM NaAc (pH 6.0). The reaction mixtures were incubated for 1 h at room temperature in the dark. Proteins were then precipitated by the addition of four volumes of acetone, followed by centrifugation at 15,000 rpm for 5 min at 4 °C. After removal of acetone, the protein pellets were air-dried and resuspended in 2× SDS gel loading buffer, followed by heating at 98 °C for 5 min. Samples were subsequently loaded onto 12% SDS–PAGE gels and electrophoresed at 180 V using Invitrogen Novex vertical gel tanks. PageRuler™ Prestained Protein Ladder (10–180 kDa; 26616, Thermo Scientific™) was used as the molecular weight marker. Fluorescent signals were detected using a Typhoon FLA 9000 imaging system (GE Healthcare Life Sciences) with the Cy3 scanning channel.

### Tyrosinase activity assay

Total proteins were extracted from leaves of WT and *ppo* mutant plants using PBS buffer. Tyrosinase activity was subsequently measured using the Tyrosinase Activity Assay Kit (Colorimetric) (ab252899, Abcam) according to the manufacturer’s instructions. Briefly, 25 μl of total protein extract was mixed with 60 μl of Tyrosinase Assay Buffer, 10 μl of Tyrosinase Substrate, and 5 μl of Tyrosinase Probe. The reaction mixtures were incubated in a microplate reader prewarmed to 37 °C, and absorbance was measured at 510 nm.

### P69B-His purification

Total protein was extracted from agroinfiltrated leaves expressing P69B-His using phosphate-buffered saline (PBS). The P69B-His protein was incubated with Ni-NTA Agarose (30210, Qiagen) at 4 °C overnight, washed three times with wash buffer (50 mM Tris-HCl, pH 7.5, 150 mM NaCl, 50 mM imidazole), and subsequently eluted using elution buffer (50 mM Tris-HCl, pH 7.5, 150 mM NaCl, 250 mM imidazole) to recover the purified protein.

### Sample preparation for Mass Spectrometry

After expressing the target protein in *Nicotiana benthamiana* leaves were collected (1g leave in 4ml PBS). After grinding in liquid nitrogen, proteins were extracted in PBS, and centrifuged to retain the input samples/supernatant (samples KZ01–KZ06, 150 μl). Part of the extract (3ml) was applied to a first round of enrichment using Ni-NTA Agarose in PBS. After three washes with Wash Buffer (50 mM pH 7.5 Tris-HCl, 150 mM NaCl, 10 mM imidazole, 0.2% Tween-20) bound proteins were eluted with Elution buffer (50 mM Tris-HCl pH7.5, 150 mM NaCl, 200 mM imidazole). The buffer was then exchanged to PBS using an Illustra™ NAP™-5 column. A 100 μl of this 1.2 ml elution was saved and precipitated with 4 Vol Acetone (samples KZ07–KZ12). Next, we performed a second round of 6xHis protein enrichment with the remaining samples. After washing three times with Wash Buffer proteins were eluted with Elution Buffer and again desalted into PBS giving samples KZ13–KZ18. All protein samples were then precipitated by addition of 4 Vol Acetone and incubation at −20 °C for 1h. The precipitated proteins were then collected by centrifugation. The supernatant was removed and the pellet dried on air briefly. Next samples were taken up in 100 µL of 1× SP3 loading buffer (1% SDS, 10 mM TCEP, 40 mM CAM, 50 mM HEPES) and heated to 90 °C for 5 min while shaking at 1300 rpm on a Thermocycler C (Eppendorf). The samples were then cleared by centrifugation and the protein concentration measured using the Pierce 660nm Assay. Subsequently we removed 15 µg of the input sample, or all of the first enrichment and 10 µg of the second enrichment and performed tryptic digestion following the SP3 sample preparation protocol (Hughes et al., 2019). Tryptic peptides were finally desalted on homemade stagetips (Rappsilber et al., 2007).

### LC-MS/MS settings

MS Experiments were performed on an Orbitrap Fusion Lumos instrument (Thermo) coupled to a Vanquish Neo ultra-performance liquid chromatography (UPLC) system (Thermo). The UPLC was operated in the one-column mode. The analytical column was a fused silica capillary (75 µm × 28 cm) with an integrated fritted emitter (CoAnn Technologies) packed in-house with Kinetex 1.7 µm core shell beads (Phenomenex). The analytical column was encased by a column oven (Sonation PRSO-V2) and attached to a nanospray flex ion source (Thermo). The column oven temperature was set to 40 °C during sample loading and data acquisition. The LC was equipped with two mobile phases: solvent A (0.2% FA, 2% Acetonitrile, ACN, 97.8% H_2_O) and solvent B (0.2% FA, 80% ACN, 19.8% H_2_O). All solvents were of UPLC grade (Honeywell). Peptides were directly loaded onto the analytical column with a maximum flow rate that would not exceed the set pressure limit of 950 bar (usually around 0.4 – 0.6 µl/min). Peptides were subsequently separated on the analytical column by running a 60 min gradient of solvent A and solvent B (for details about gradient composition, see **Supplemental Tables S3-S7**) at a flow rate of 300 nl/min. The mass spectrometer was controlled by the Orbitrap Fusion Lumos Tune Application (version 4.1.4244) and operated using the Xcalibur software (version 4.7.69.37). Detailed settings for the mass spectrometer can be found in **Supplemental Tables S3-S7**.

### Data Processing Protocol CL-MS

RAW spectra were submitted to a closed MSFragger (version 4.1, Kong et al., 2017) search in Fragpipe (version 22) (Yu et al., 2023) using the “LFQ-MBR” workflow (label-free quantification and match-between-runs; default settings were used unless otherwise stated). RAW files were listed in the “Input LC-MS Files” section and experiment set “by file name”. As “Data Type” we kept the default “DDA” (data dependent acquisition). The MS/MS spectra were searched against a custom database generated in Fragpipe 2025-11-14-decoys-contam-ACE_1035_NbLab360.v103.gff3.CDS.fasta.AA_plus_SOI.fasta.fas (45850 entries including contaminants and the same number of decoys). The database comprises mainly of the *N. benthamiana* reference proteome NbLab360.v103.gff3.CDS.fasta (45730 entries) plus the two sequences of interest (P69B and EPI1). MSFragger searches allowed oxidation of methionine residues (16 Da; 3) and acetylation of the protein N-terminus (42 Da; 1), as variable modifications (first value in brackets refers to the molecular weight of the modification, second value to the maximum number of occurrences per peptide). A maximum of 3 variable modifications and a maximum of 5 combinations were allowed globally. Carbamidomethylation on cysteine (57) was selected as static modification. Enzyme specificity was set to “Trypsin”. The initial precursor and fragment mass tolerance was kept at ±20 ppm. Mass calibration and parameter optimization was selected. Validation of peptide spectrum matches was done by running MSBooster using DIA-NN for RT and spectra prediction. Peptide spectra matches (PSM) were validated using percolator with a minimum probability setting of 0.5. Protein inference was performed using ProteinProphet (part of Philosopher version 5.1.1). The final reported protein FDR was 0.01 (based on target-decoy approach). Protein quantification was performed with IonQuant (version 1.10.27). MaxLFQ (minions 1), MBR (FDR 0.01) and normalization of intensities across runs was selected. Unique and razor peptides were allowed. Advanced options were kept at default. Further analysis and filtering of the results were done in Perseus v1.6.10.0. Comparison of protein group quantities (relative quantification) between different MS runs is based solely on the LFQs as calculated by IonQuant, MaxLFQ algorithm.

### Statistical analysis

All statistical analyses were performed in GraphPad Prism (version 11.0.0). Comparisons between WT and mutant lines were made using one-way ANOVA (ordinary, Welch’s, or Kruskal–Wallis, depending on whether assumptions of normality and homoscedasticity were met), with post hoc comparisons versus WT conducted using Dunnett’s, Dunnett’s T3, or Dunn’s test as appropriate. For experiments involving multiple factors, two-way ANOVA was performed with Geisser–Greenhouse correction applied where relevant, and post hoc comparisons versus WT used Dunnett’s or Šídák-adjusted tests. Data are presented as mean ± SEM, and *P* < 0.05 was considered statistically significant.

## Results

### Generation and characterization of *ppo* double knockout lines

The 300bp fragment that we previously used to silence *PPO* (Mahadevan et al., 2025) targets two highly homologous *PPO* genes: *PPO1* (NbL17g16540) and *PPO2* (NbL10g18390). According to LAB360 and NbenBase genome annotations (Ranawaka et al., 2023; Kurotani et al. 2023), *Nicotiana benthamiana* has seven *PPO* genes (**Supplemental Fig. S1a**). However, only *PPO1* is significantly expressed in leaves, and its expression increases upon agroinfiltration (Grosse-Holz et al., 2018, Supplemental **Fig. S1b**). To generate the *ppo* knockout lines, we designed two single guide RNAs (sgRNAs) targeting these *PPO* genes and we selected two independent homozygous lines carrying disrupted open reading frames for both genes. Line #1 carries a 284bp deletion in *PPO1* (allele *ppo1-1*), and 2bp and 1bp deletions in *PPO2* (allele *ppo2-1*), whereas line #2 carries a different 284bp deletion in *PPO1* (allele *ppo1-2*) and a 281bp inversion in PPO2 (allele *ppo2-2*) (**Fig. 1a**). Consequently, both *PPO1* and *PPO2* were disrupted early in the open reading frames before encoding the catalytic residues (**Fig. 1b**), indicating that these lines are double null mutants for both *PPO* genes. Western blot analysis of leaf extracts with anti-PPO antibodies indeed confirmed that both lines lack the PPO protein (**Fig. 1c**). A Ponceau-S stain of the corresponding membrane shows equal loading but also displays a 180 kDa high molecular weight (HMW) signal for WT plants that is absent from *ppo* mutant lines (**Fig. 1h**).

**Fig. 1.**
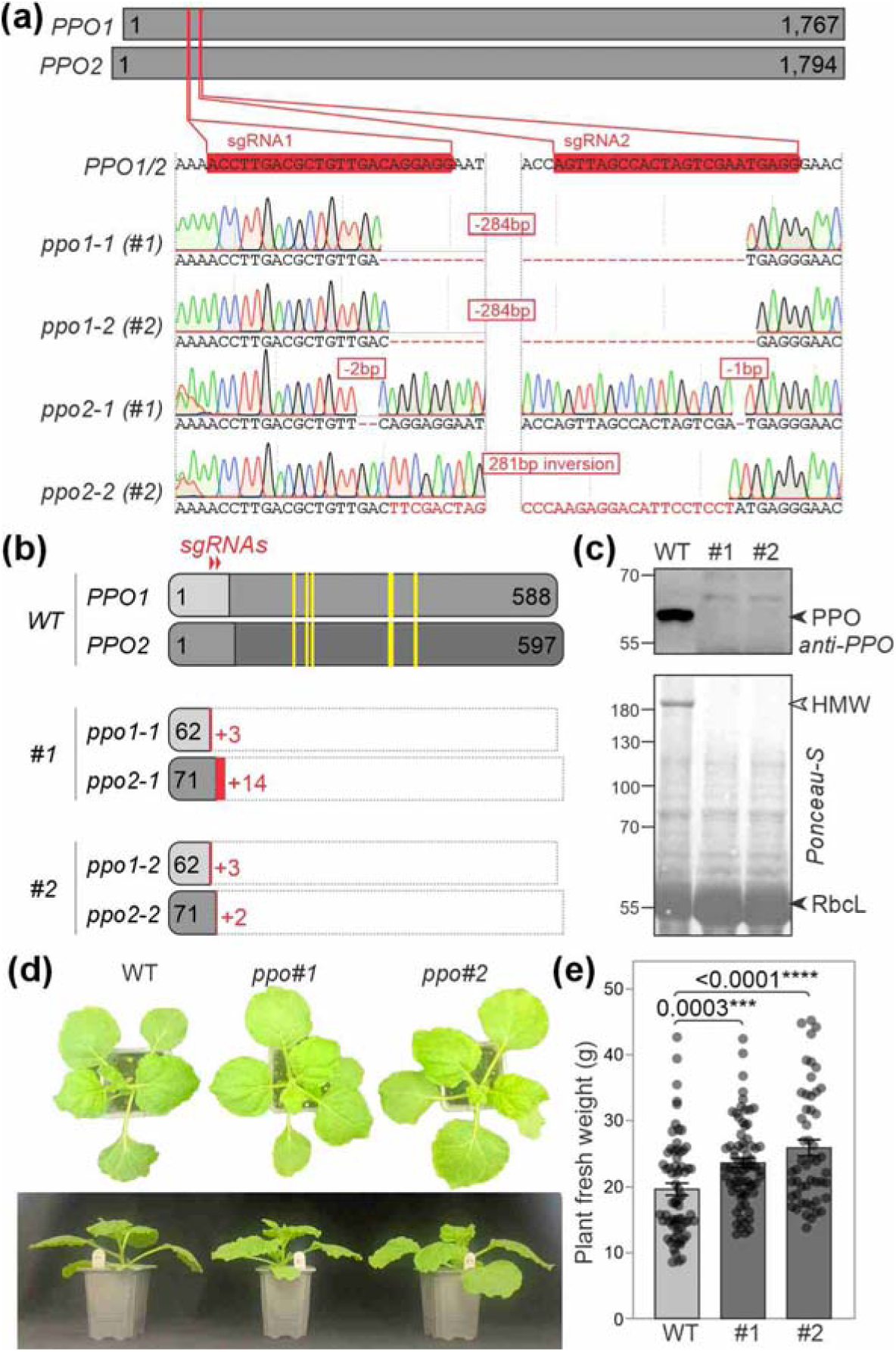
Characteristics of *ppo* double mutant knockout lines. **(a)** CRISPR/Cas9-induced nucleotide alterations in *PPO* genes. The 5’ region of the open reading frames of *PPO1* and *PPO2* were targeted by two single guide RNAs (sgRNAs, red). The sgRNA target locus were sequenced in homozygous lines #1 and #2 carrying alleles *ppo1-1* and *ppo2-1* in line #1 or alleles *ppo1-2* and *ppo2-2* in line #2, respectively. Shown are the alignments with wild-type *PPO1* and *PPO2*, which are identical in the shown regions. **(b)** Predicted proteins encoded by the mutant *ppo* alleles. The length of the open reading frame including the nonsense sequence (red) is indicated in amino acid residues. The prodomain (light grey) and catalytic residues (yellow) are highlighted. **(c)** The PPO protein is absent in leaf extracts of *ppo* mutants. Leaf extracts from 4-week-old plants were analysed by Ponceau-S staining and western blot analysis using the anti-PPO antibody. **(d)** Top image of 28-day old WT and *ppo* mutant plants. **(e)** Mutant *ppo* plants are slightly larger than wild-type plants. The fresh weight of the plants above ground was measured in four sets of 30-31 days old plants. Error bars represent standard error of n>50 replicates generated in four batches of plants. Statistical significance was assessed using ANOVA.

Both *ppo* mutants grew and developed normally (**Fig. 1d**), but the *ppo* mutants seemed to grow slightly larger. The *ppo* mutants indeed accumulate 20-32% more fresh weight (**Fig. 1e**), which is a great benefit to protein production yields.

### PPO depletion reduces enzymatic browning and native protein crosslinking

PPO accumulation clearly correlates with leaf age, with most PPO protein accumulating in young, expanding leaves, and being absent in both *ppo* mutants (**Fig. 2a**). However, the 180 kDa HMW signal is strongest in mature leaves (**Fig. 2a**), which are most often selected for agroinfiltration for their high expression and ease of infiltration.

**Fig. 2.**
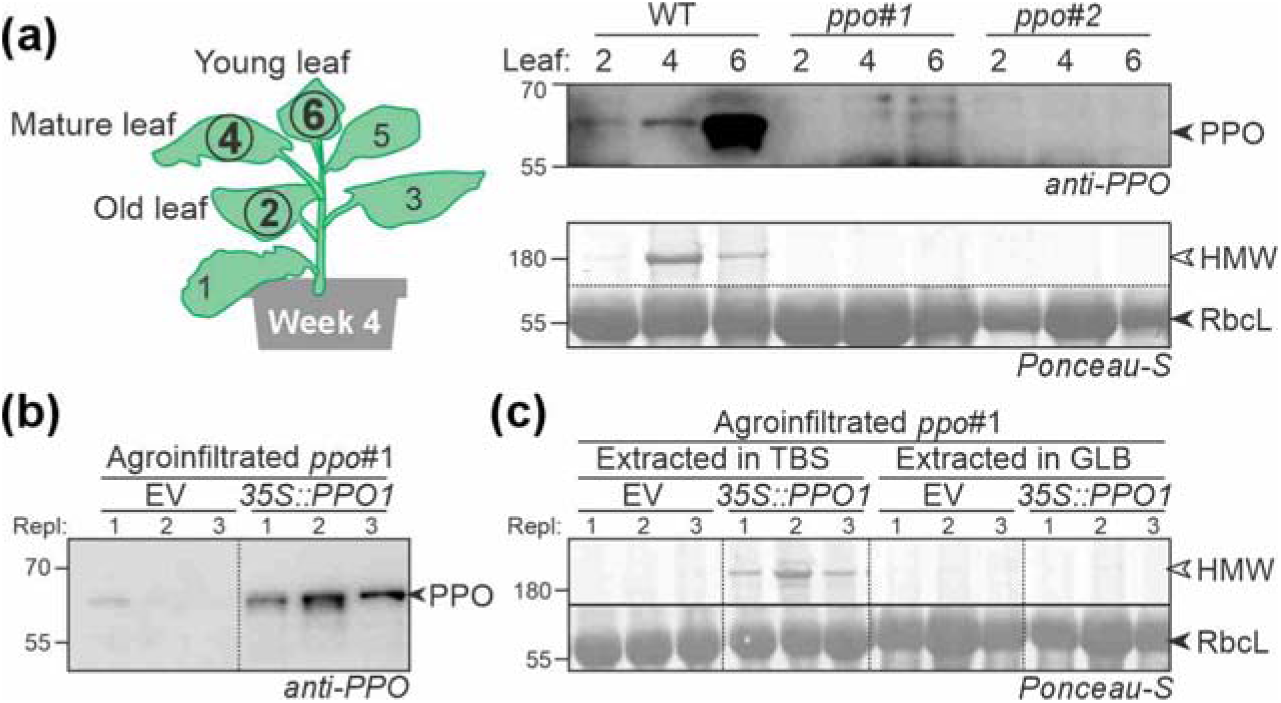
PPO is responsible for browning and crosslinking. **(a)** Native crosslinking into high molecular weight (HMW) complexes in mature leaves is absent in *ppo* mutants. Leaf extracts from leaf discs of young, mature and old leaves were generated in TBS and analysed by Ponceau-S staining and western blot analysis using the anti-PPO antibody. **(b)** Transient expression of *35S::PPO1* triggers PPO accumulation in *ppo* mutant. Leaves of the *ppo* mutant #1 were agroinfiltrated and samples were analysed 4 days later by western blot using the anti-PPO antibody. **(c)** Native crosslinking into the HMW complex occurs in extracts of PPO1 expressing leaves in Tris-buffered saline (TBS), but not in gel-loading buffer (GLB).

To confirm the role of PPO in native crosslinking, we transiently overexpressed PPO1 and the empty vector (EV) in the *ppo* mutant by agroinfiltration. Western blot analysis confirmed that PPO accumulated in leaves agroinfiltrated with *35S::PPO1* and not in the EV control (**Fig. 2b**), but PPO levels are not much higher than endogenous PPO levels, despite the use of the strong CaMV 35S promoter. Ponceau-S staining revealed that the 180 kDa HMW signal accumulated upon PPO expression when extracts were generated in TBS, but not upon extraction in gel loading buffer (GLB) containing reducing agent dithiothreitol (DTT) and sodium dodecyl sulphate (SDS) (**Fig. 2c**), confirming that PPO-induced protein crosslinking occurs during extraction and that PPO inactivation can prevent this.

### Transient protein expression is not altered in *ppo* mutants

We previously noticed that transient expression of GFP might be higher in *TRV::PPO* plants (Mahadevan et al., 2025). In our *ppo* mutants, however, fluorescence is not increased upon transient GFP expression (**Supplemental Fig. S2a**), and similar levels of GFP protein accumulated in the *ppo* mutants (**Supplemental Fig. S2b**), indicating that transient expression of GFP is not significantly different in *ppo* mutants compared to WT plants.

### PPO depletion preserves endogenous proteins at predicted molecular weights

To investigate how PPO depletion affects the integrity of endogenous proteins under non-denaturing extraction conditions, we incubated total leaf extracts from wild-type and *ppo* mutant plants in phosphate-buffered saline for one hour and analysed protein migration by SDS–PAGE. Coomassie staining revealed that incubation of wild-type extracts resulted in a pronounced loss of proteins migrating at their predicted molecular weights, accompanied by the appearance of a prominent high-molecular-weight (HMW) signal around 180–200 kDa (**Fig. 3a**). In contrast, extracts from *ppo* mutants retained substantially stronger signals at their expected molecular weights and displayed a marked reduction in HMW material (**Fig. 3a**).

**Fig. 3.**
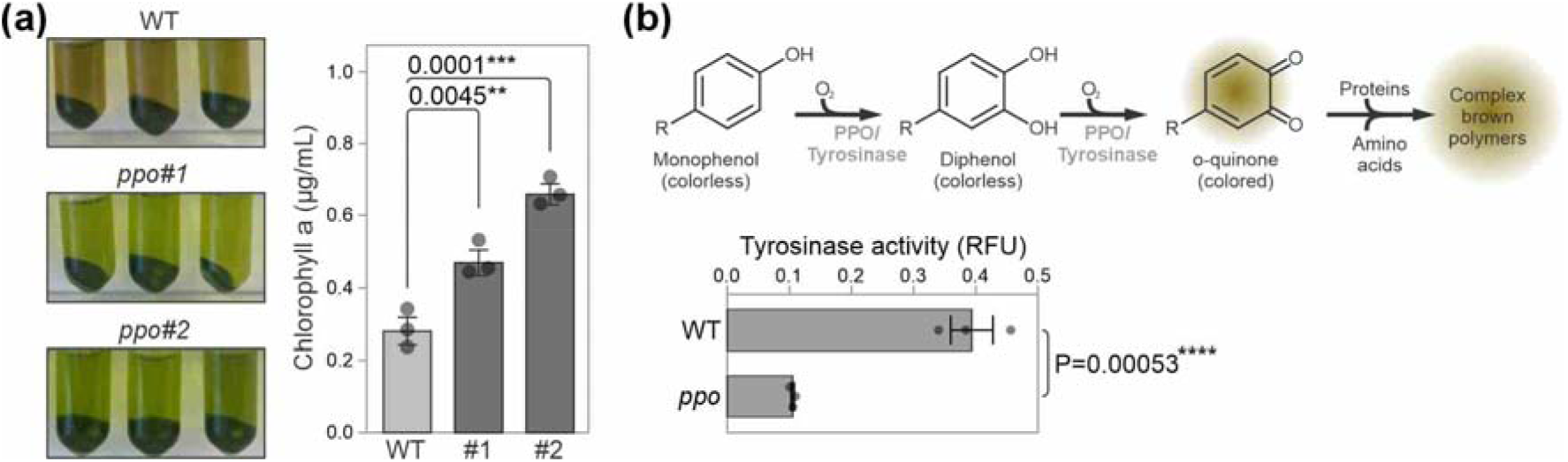
PPO depletion preserves endogenous proteins at their predicted molecular weights and enhances detectable enzymatic activities. **(a)** Less RbcL crosslinking in *ppo* mutant. Leaf extracts were incubated for 1h, separated on protein gels, stained with Coomassie or analysed by Western blot with the anti-RbcL antibody on n=3 replicates. **(b)** Increased levels of endogenous enzymes at their predicted molecular weight. Proteomes used in (A) were probed with antibodies against fructose biphosphate aldolase (ALD) and serine hydroxymethyl transferase (SHMT). **(c)** More activity-based signals detected in extracts of *ppo* mutants. Leaf extracts were labeled with FP-TAMRA, which covalently binds active serine hydrolases for 60 minutes, separated on protein gels and scanned for fluorescence in n=3 replicates.

Immunoblot analysis using an antibody against RbcL demonstrated that the HMW signal observed in wild-type extracts contains RbcL, whereas RbcL in *ppo* mutant extracts predominantly migrated at its predicted molecular weight of approximately 55 kDa (**Fig. 3a**). Quantification revealed that the RbcL signal at 55 kDa was approximately twofold higher in *ppo* mutant extracts compared to wild type, indicating that PPO depletion preserves RbcL in its native, non-crosslinked form.

To assess whether this effect extends beyond RbcL, we probed the same extracts with antibodies against fructose-bisphosphate aldolase (ALD) and serine hydroxymethyltransferase (SHMT), two abundant metabolic enzymes with predicted molecular weights of 41.9 kDa and 54.8 kDa, respectively. Both ALD and SHMT displayed significantly stronger signals at their predicted molecular weights in *ppo* mutant extracts compared to wild type, with increases of 2.32-fold and 1.48-fold, respectively (**Fig. 3b**). These results indicate that PPO-mediated crosslinking broadly affects multiple endogenous proteins during extraction and that genetic removal of PPO preserves a wider range of proteins in their native molecular forms.

Because preservation of protein integrity is expected to improve the detection of enzymatic activities, we next performed activity-based profiling of serine hydrolases using the fluorescent probe FP-TAMRA. FP-TAMRA is a fluorescent fluoro-phosphonate probe that reacts with hyperreactive serine residues in the active site of serine hydrolases in an activity-dependent manner (Liu et al., 1999). Labeling of extracts from *ppo* mutant plants revealed 9 additional fluorescent signals compared to wild-type extracts (**Fig. 3c**). These additional signals indicate that more active serine hydrolases are detectable in the absence of PPO-mediated crosslinking, suggesting that PPO activity limits the measurable activity landscape of endogenous enzymes in leaf extracts.

### PPO-dependent tyrosinase activity underlies native protein crosslinking

Because the leaf extracts of *ppo* mutants not only appeared less brown, but also much greener (**Fig. 4a**), we quantified chlorophyll levels in acetone extracts by absorbance. These measurements indicate a significant 68-135% increase of chlorophyll-a in *ppo* mutants (**Fig. 4a**), suggesting that chlorophyll is oxidized by PPO in leaf extracts.

**Fig. 4.**
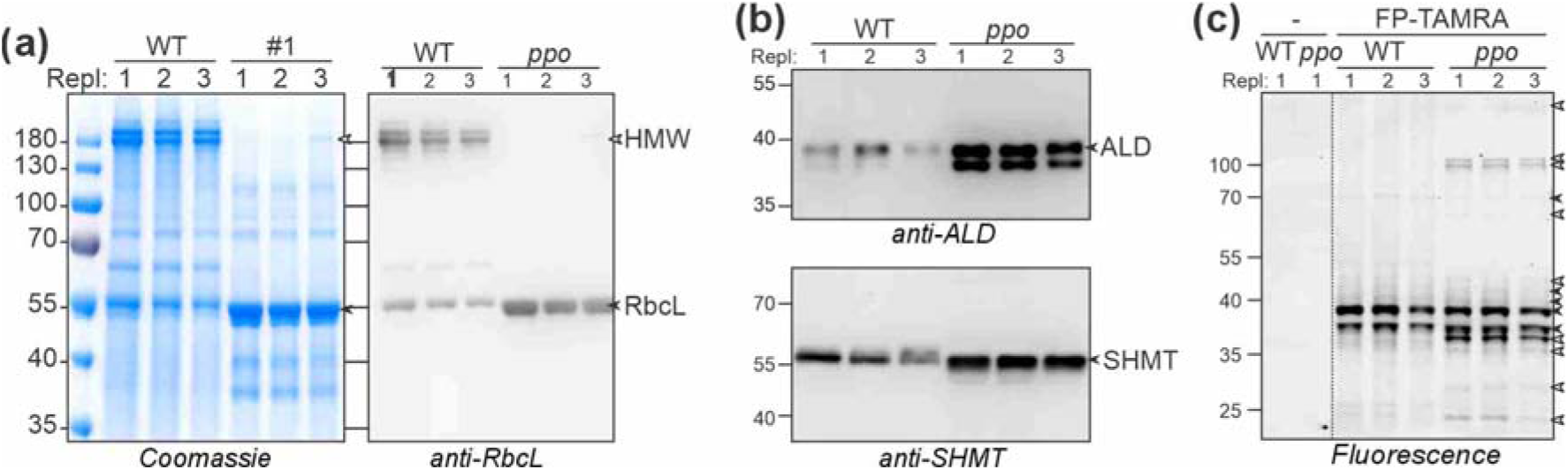
Polyphenol oxidase depletion reduces enzymatic browning and tyrosinase activity in *Nicotiana benthamiana* leaf extracts. **(a)** Leaf extracts of *ppo* mutants are less brown and greener and contain higher chlorophyll-a levels. Left: leaf extracts were incubated in TBS for 2 min and centrifugated before the image was taken. Right: leaf extracts were incubated with 80% acetone and the concentrations of Chlorophyll-a were measured by absorbance spectroscopy. Error bars represent standard error of n=3 biological replicates. P-values were calculated with ANOVA. **(b)** Phenol oxidation by PPO/Tyrosinase can cause protein polymers (upper panel). Reduced tyrosinase activity in extracts of *ppo* mutants (lower panel). Leaf extracts were assayed for tyrosinase activity in PBS. Error bars represent SE of n=3 replicates. P-value was calculated with Students t-test.

Polyphenol oxidases can promote protein crosslinking through two related mechanisms: the oxidation of phenolic compounds released during tissue disruption and the direct oxidation of solvent-exposed tyrosine residues on proteins via tyrosinase activity. Both processes generate reactive quinones that can form covalent bonds with nucleophilic amino acid side chains, resulting in protein polymerization (**Fig. 4b**).

To determine whether PPO-dependent tyrosinase activity contributes to the observed native protein crosslinking, we measured tyrosinase activity in total leaf extracts from wild-type and *ppo* mutant plants. Extracts from *ppo* mutants displayed a threefold reduction in tyrosinase activity compared to wild type (**Fig. 4b**). This reduction is consistent with the absence of PPO protein in the mutant lines and indicates that PPO is the major contributor to tyrosinase activity in leaf extracts under the tested conditions.

These results support a model in which PPO-mediated oxidation reactions, including tyrosine oxidation, drive the formation of high-molecular-weight protein complexes during extraction. The strong reduction of both tyrosinase activity and protein crosslinking in *ppo* mutants demonstrates that genetic removal of PPO effectively suppresses these processes.

### Recombinant protein purification yield and purity are improved in *ppo* mutants

Native protein crosslinking during extraction is expected to reduce the recovery of recombinant proteins by trapping them in insoluble or high-molecular-weight complexes. To test whether PPO depletion improves recombinant protein purification, we transiently expressed a His-tagged version of the tomato subtilase P69B in wild-type and *ppo* mutant plants. P69B is defence-related subtilase of tomato that we study as a target for various pathogen-secreted proteins (Homma et al., 2023). Immunoblot analysis of total leaf extracts revealed comparable levels of P69B-His accumulation in both genotypes, indicating that PPO depletion does not affect transient expression of this recombinant protein (**Supplemental Fig. S3**).

Following extraction under non-denaturing conditions, P69B-His was purified using Ni–NTA affinity chromatography. While similar amounts of P69B-His were captured on the affinity resin from wild-type and *ppo* mutant extracts, the amount of P69B-His recovered in the eluate was significantly higher when purified from *ppo* mutant plants (**Fig. 5a**). Coomassie staining revealed the HMW signal is present in the eluate of WT plants, but much less in that of *ppo* mutant plants (**Fig. 5a**), indicating that eluates from *ppo* plants are much cleaner when compared to those from WT plants.

**Fig. 5.**
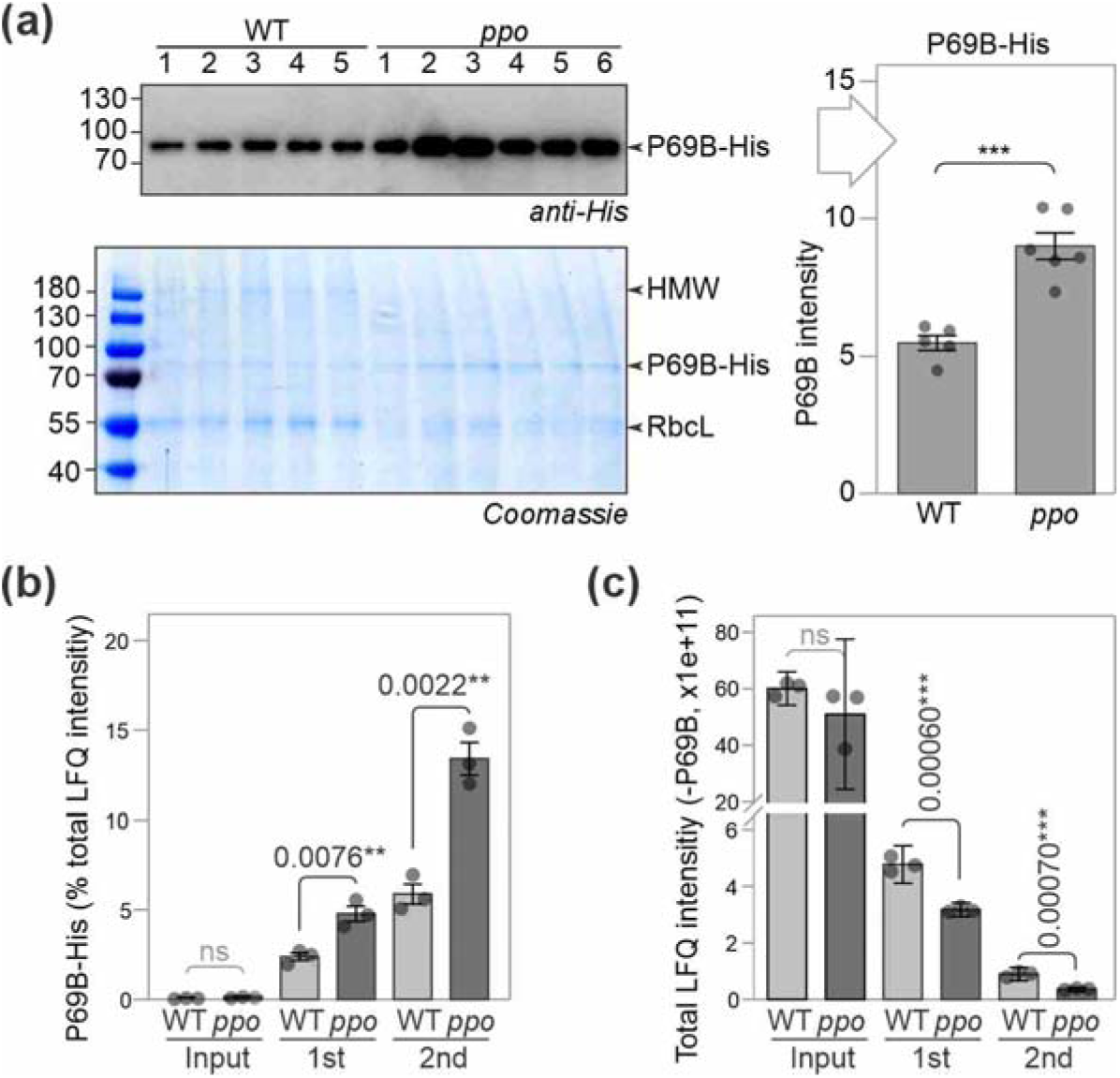
PPO depletion improves the yield and purity of a recombinant protein purified from *Nicotiana benthamiana*. **(a)** More P69B-His was purified from the *ppo* mutant. Leaf total protein extracts (Extract) of WT and *ppo* mutant line #1 transiently expressing P69B-His at 3dpi were passed over Ni-NTA beads in n=5-6 replicates. Beads were washed and the imidazole-eluted fraction samples were analysed by western blot using the P69B-antibody and Coomassie staining. Western blot signals were quantified and plotted. Error bars represent SE of n=5 replicates. P-value: Student’s t-test, < 0.001 (***). **(b)** Increased purity of P69B from the *ppo* mutant. P69B-His was purified twice over Ni-NTA from extracts of leaves transiently expressing P69B and input and eluate samples were analysed by MS for n=3 replicates. Ion intensities of P69B-His determined by label-free quantification (LFQ) were calculated as a percentage of the total LFQ intensity. Error bars represent SE of n=3 replicates. P-values: Student’s t-test. **(c)** Reduced intensity of contaminant proteins from *ppo* mutant. Total LFQ intensities without P69B-His are shown for samples described in (b). Error bars represent SE of n=3 replicates. P-values: Student’s t-test.

To further quantify purification efficiency and purity, we performed mass spectrometry analysis on independently purified P69B-His samples, including a second round of Ni–NTA enrichment. The purity of P69B-His was more than 2-fold higher from extracts of *ppo* mutants after both purifications, reaching 14% of the total ion intensities (**Fig. 5b**). The increased purity corresponded with significantly lower ion intensities of protein contaminants in the *ppo* mutant (**Fig. 5c**).

Together, these results demonstrate that PPO depletion substantially improves both the yield and purity of recombinant proteins purified from *N. benthamiana* leaf extracts, without altering their expression levels.

## Discussion

### PPO-mediated crosslinking represents a bottleneck in plant protein extraction

This study demonstrates that PPOs impose a substantial and previously underappreciated constraint on protein integrity during extraction from *Nicotiana benthamiana* leaves. Genetic disruption of the two major leaf-expressed *PPO* genes resulted in viable *ppo* double knockout plants without obvious developmental defects and with slightly increased biomass (**Figs. 1d and 1e**), indicating that PPO depletion is well tolerated under standard growth conditions. Despite this mild whole-plant phenotype, PPO removal had profound consequences for protein integrity during extraction.

While enzymatic browning has long been recognized as a nuisance in plant protein purification, our results show that PPO activity has broader consequences by driving extensive native protein crosslinking that alters protein migration, reduces detectable enzymatic activities, and compromises recombinant protein recovery (**Figs. 2–5**). In wild-type plants, extraction under non-denaturing conditions resulted in rapid browning and the formation of high-molecular-weight protein material (**Figs. 2b and 4a**), illustrating how quickly PPO-mediated reactions are triggered upon tissue disruption.

At the protein level, PPO-mediated oxidation caused abundant stromal enzymes such as RbcL to shift into high-molecular-weight complexes in wild-type extracts, whereas RbcL remained predominantly at its predicted molecular weight in *ppo* mutant extracts (**Fig. 3a**). This effect was not restricted to RbcL but extended to other endogenous enzymes, including fructose-bisphosphate aldolase and serine hydroxymethyltransferase (**Fig. 3b**), indicating that PPO-mediated crosslinking broadly affects the leaf proteome.

### Consequences of PPO depletion for endogenous enzyme activity profiling

The preservation of proteins at their predicted molecular weights in *ppo* mutants was accompanied by an increase in detectable enzymatic activities. Activity-based protein profiling revealed a greater number of active serine hydrolases in extracts from *ppo* mutants compared with wild type (**Fig. 3c**), indicating that PPO activity restricts the measurable activity landscape of endogenous enzymes. Because activity-based probes require access to intact and catalytically competent active sites, these results suggest that PPO-mediated oxidation and crosslinking mask or inactivate enzyme activities in wild-type extracts. Taken together, our results indicate that *ppo* mutants will offer significant advantages in studies on native proteomes, including protein activities and protein-protein interactions.

Mechanistically, PPOs contribute to the oxidative capacity of leaf extracts through both phenolic substrates and oxidation of solvent-exposed tyrosine residues. Consistent with this, extracts from *ppo* mutants displayed strongly reduced tyrosinase activity (**Fig. 4b**). Together with the conceptual model shown in **Fig. 4b**, these data support a mechanism in which PPO activity increases the oxidative potential for protein browning and polymer formation during extraction, thereby indirectly limiting enzyme activity detection.

These observations have broader implications for functional proteomics in plants. Many studies aim to characterize enzyme activities under near-native conditions, yet the contribution of PPO-mediated reactions to activity loss has not been systematically considered. Our results indicate that PPO depletion provides a straightforward genetic strategy to improve the reliability of activity-based profiling and biochemical assays performed on leaf extracts.

### Implications for recombinant protein yield and purity in molecular pharming

Recombinant protein production in *N. benthamiana* relies heavily on downstream purification from total leaf extracts, where native crosslinking can trap proteins in insoluble or heterogeneous assemblies. Importantly, PPO depletion did not increase transient expression levels, as GFP accumulation and fluorescence were comparable between wild-type and *ppo* mutant plants (**Supplemental Figs. S2a and S2b**). This demonstrates that differences observed during purification cannot be attributed to altered expression efficiency.

Instead, PPO depletion substantially improved downstream recovery. The significant increase in P69B-His recovery from *ppo* mutant extracts (**Fig. 5a**) demonstrates that PPO-mediated crosslinking constitutes a major bottleneck during purification. Moreover, mass spectrometry–based analysis revealed a marked increase in purity of recombinant protein preparations derived from *ppo* mutants, accompanied by a corresponding reduction in contaminating plant proteins (**Figs. 5b and 5c**).

These findings extend previous observations made using transient *PPO* silencing approaches and demonstrate that stable genome-edited *PPO* knockout lines provide consistent and robust benefits for protein purification (Mahadevan et al., 2020). Unlike chemical inhibitors or transient silencing strategies, genetic removal of *PPO* eliminates the need for additional treatments and avoids variability associated with incomplete suppression.

Our findings are consistent with a recent report that knocking out the same two *PPOs* from *Nicotiana benthamiana* improves recombinant protein purification (Diao et al., 2026). Similar to our observations, Diao and colleagues found no increased transient expression of GFP in *ppo* mutants, and reported significant improvements in yield and purity of recombinant proteins upon purification.

### Comparison with alternative strategies to mitigate extraction-induced artifacts

Various strategies have been employed to mitigate enzymatic browning and protein degradation during plant protein extraction, including the use of reducing agents, polyvinylpolypyrrolidone, protease inhibitors, and rapid denaturation. While such approaches can partially suppress browning, they are often incompatible with experiments requiring native protein structure and activity. In contrast, genetic removal of *PPO* intrinsically suppresses a major source of oxidation-driven crosslinking at its origin, as evidenced by reduced browning, preserved protein migration, and improved enzyme activity detection (**Figs. 2–4**).

Recent advances in genome editing, VIGS and RNAi technology have facilitated the generation of plant lines optimized for transient expression and protein production, such as plants deficient in immune receptors (Dodds et al., 2025) or proteases (Beritza et al., 2024b) and alkaloids (Vollheyde et al., 2023). The *ppo* mutants described here complement these approaches by addressing a distinct but equally critical bottleneck at the extraction and purification stage.

### Future applications

PPOs have been implicated in plant defense and stress responses (Zhang and Sun 2021; Zhang 2023), and their removal may influence susceptibility to pests or pathogens under certain conditions (Felton et al., 1989; Constabel et al., 1995; Thipyapong et al., 2004; Bhonwong et al., 2009). However, the absence of obvious developmental defects and the slightly increased biomass observed in *ppo* mutants suggest that PPO deletion is well tolerated under controlled growth conditions (**Figs. 1d, 1e**).

Future studies could explore the combination of *PPO* knockout lines with other genetic modifications designed to enhance transient expression, reduce proteolysis, or suppress immune responses. Such combinatorial approaches may further optimize *N. benthamiana* as a versatile chassis for plant science and molecular pharming. In addition, extending PPO depletion strategies to other plant species used for protein production could broaden the applicability of these findings.

## Conclusion

We have introduced two independent *ppo* double mutant lines that lack the PPO protein, and have less native crosslinking and browning in leaf extracts. With reduced native crosslinking, leaf extracts of *ppo* mutants maintain native proteins at their predicted molecular weight and facilitate the detection of more enzyme activities. Transient GFP expression is not affected in *ppo* mutants, but the purification yield and purity of recombinant proteins are significantly improved. These data indicate that these *ppo* lines will be instrumental for both plant science and molecular pharming industry by enabling the production and study of purified proteins in their native state *in planta*.

## Acknowledgements

We thank Ursula Pyzio for excellent plant care; Dr. Shi-jian Song for ordering the *ppo* KO lines; and Sarah Rodgers, Caroline O’Brien, Patricia Bowman, Jenny Borman and Svenja Heimann for technical support. This project was financially supported by BBSRC project BB/Y00969X/1 (KZ), the BBSRC Interdisciplinary DTP DDT00230 (EW), and ERC-AdG-2020 project 101019324 ‘ExtraImmune’ (RH).

## Author Contributions

RH conceived the project; KZ performed the experiments; FK and MK did proteomic analysis; EW did bioinformatic analysis and statistics; RH and KZ wrote the manuscript with input from all authors.

## Data Availability Statement

The mass spectrometry proteomics data for the digestions have been deposited to the ProteomeXchange Consortium via the PRIDE (Vizcaíno et al., 2016) partner repository (https://www.ebi.ac.uk/pride/archive/) with the dataset identifier PXD071885. During the review process, the data can be accessed via a reviewer account (Username; Password: QI7GqsRycNe7).

## SUPPLEMENTAL TABLES

**Table S1.**
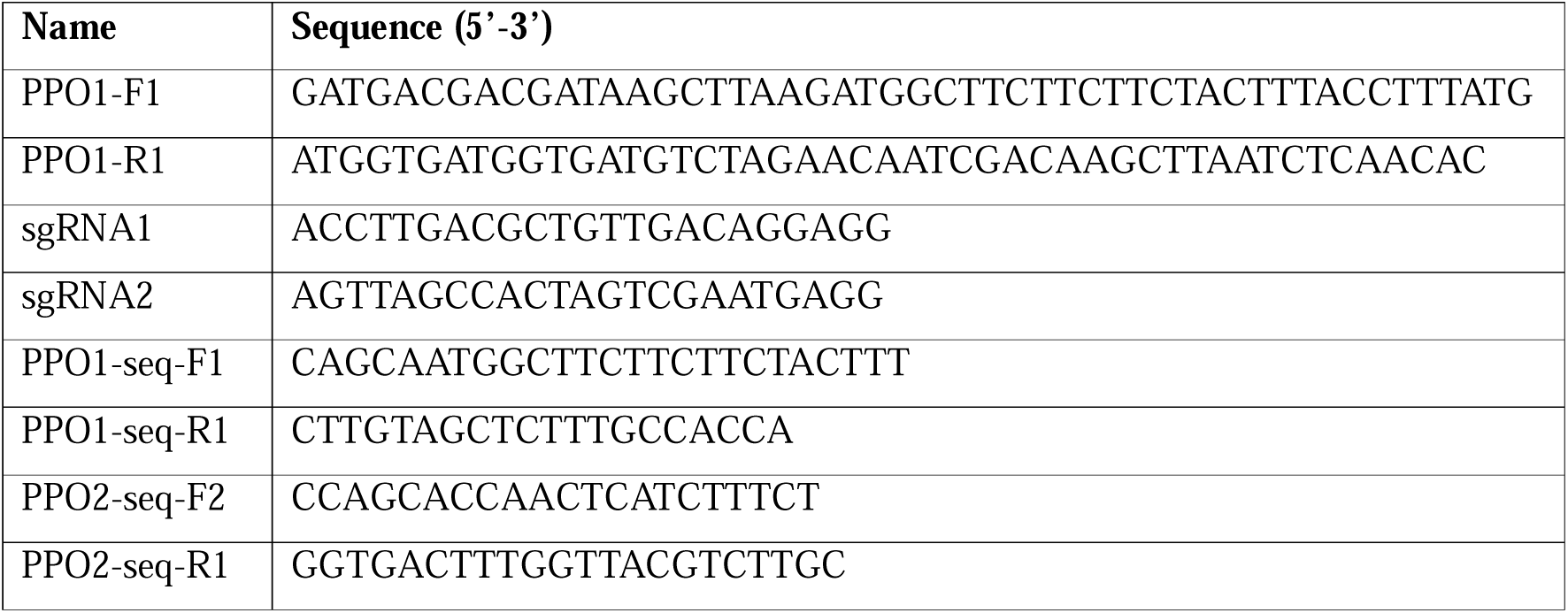
Used oligonucleotides.

**Table S2.** Used plasmids.

| Name | Description | Reference |
| --- | --- | --- |
| pFH20 | Binary vector 35S:NtPR1a-P69B-His | Homma et al., 2023 |
| pKZ165 | Binary vector 35S:nlsGFP | This work |
| pKZ205 | Binary vector 35S:FLAG-PPO1-2xHis | This work |
| pJK001c | Empty binary vector | Paulus et al., 2020 |
| P19 | Binary 35S::P19 | Van der Hoorn et al., 2003 |

---

**Table S3.**
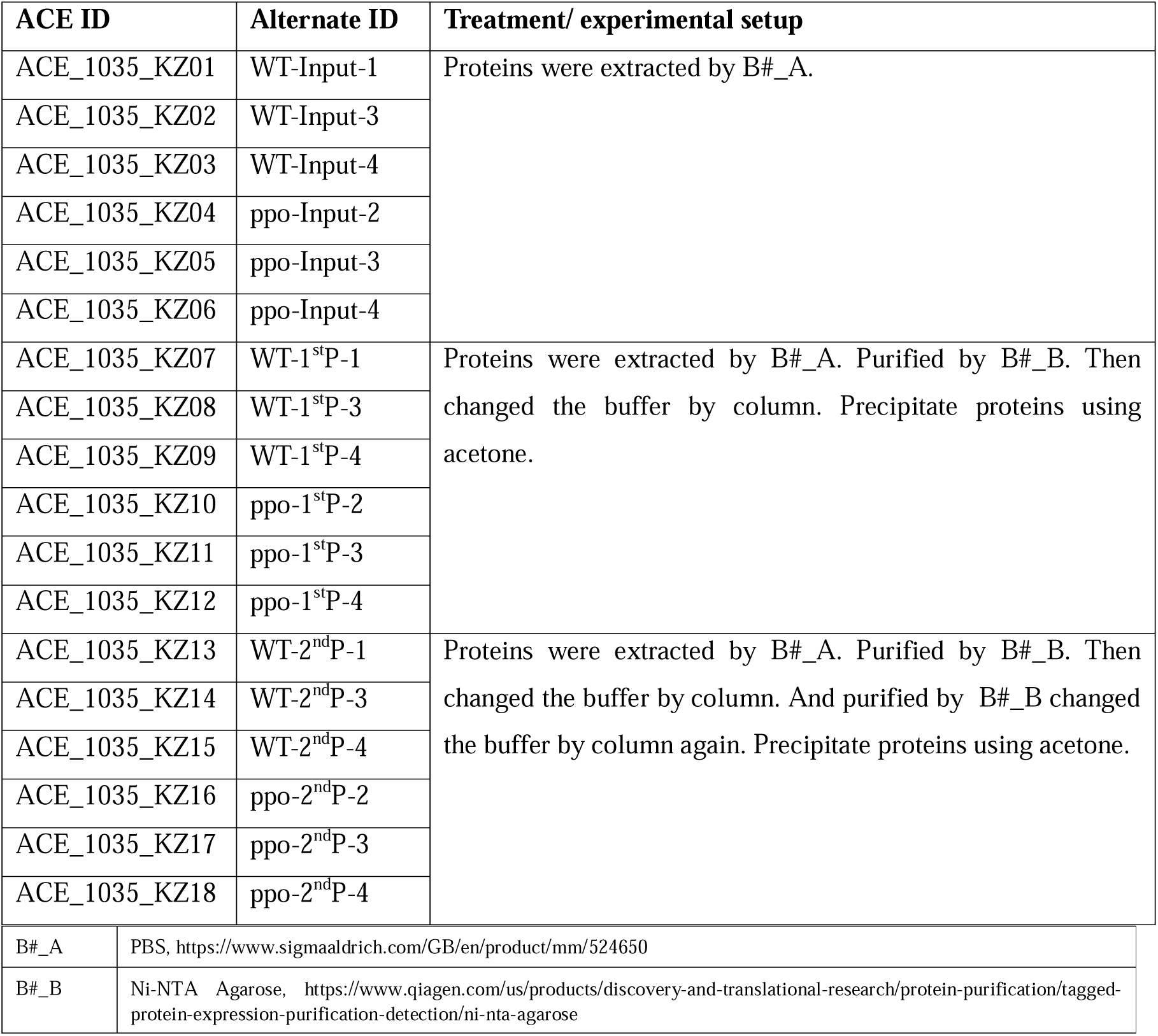
Samples used for proteomics.

**Table S4.**
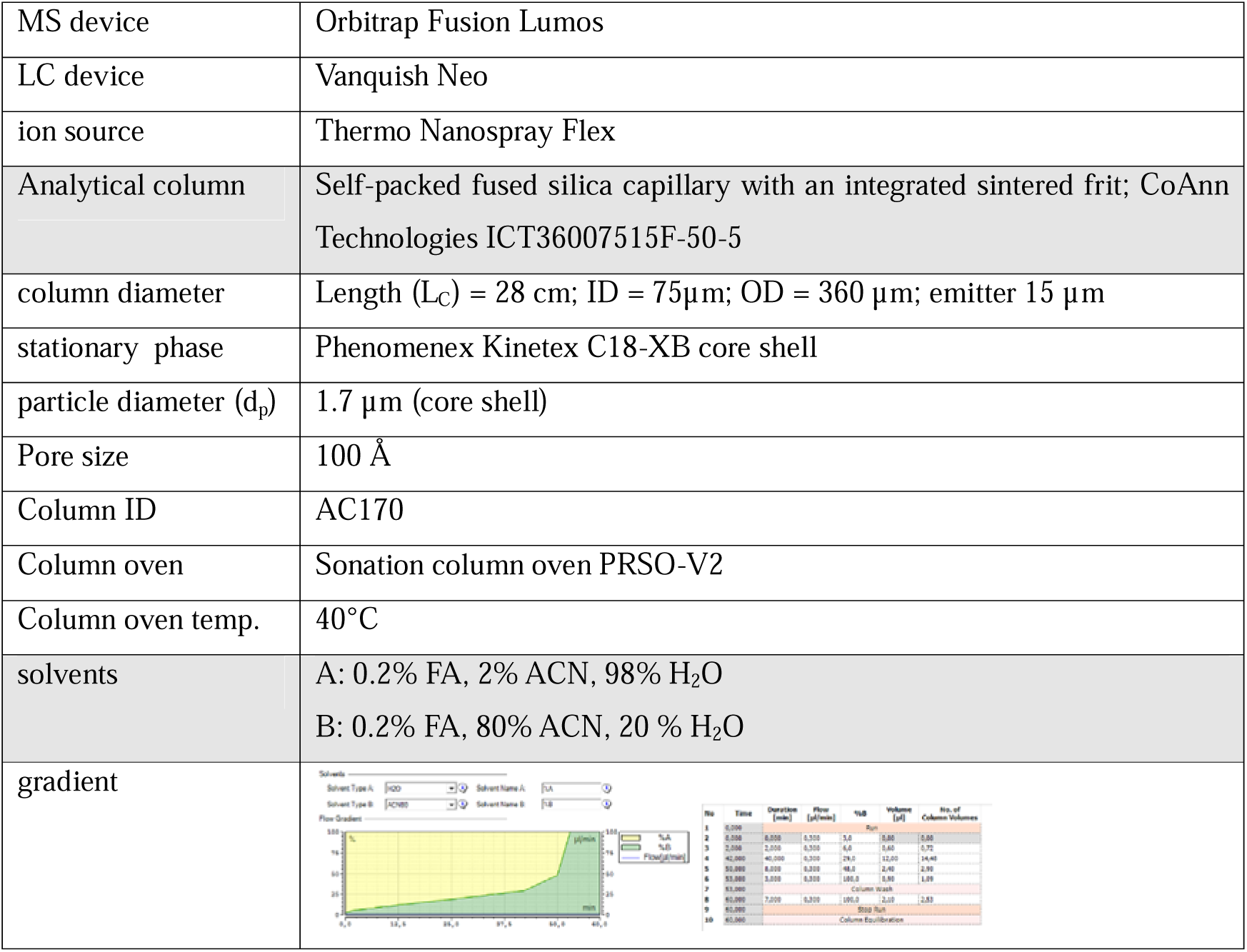
LC setting.

**Table S5.** MS settings.

| General | MS1 | MS2 | General comments; special settings |
| --- | --- | --- | --- |
| Lumos<br>Tune v4.1.4244<br>Xcalibur<br>v4.7.69.37<br>SII: 1.7.0.468<br>Gradient: 60 min | Analyzer: FT<br>Res.: 120000<br>SR: 375 - 1500<br>AGC: Standard<br>AGC abs.: 400000<br>AcT: 50 ms<br>RF: 30<br>IsM: Q<br>SF: --<br>DDM: CT/3sec | Analyzer: IT<br>Res./ScR: -/rapid<br>SR: Auto<br>AGC: 300%<br>AGC abs.: 30000<br>AcT: 35 ms<br>CS: +2 to +7<br>IsM: Q<br>IsW: 1.6<br>Frag.: HCD<br>NCE: 32 | classic orbitrap experiment: MS1 in Orbitrap at high resolution and data dependent MS2 also in Orbitrap high resolution. Dynamic exclusion enabled (exclude after n times=1; Exclusion duration (s)= 30; mass tolerance= ± 10ppm)<br>Intensity Threshold: 5000<br>Ion transfer Tube Temp: 250 °C<br>Ion Source Voltage: 2500 V |

Note: FT= Fourier Transform (Orbitrap); IT= Iontrap; Q= Quadrupol; Res.= max. Resolution at 200 m/z (Lumos) or 400 m/z (Elite) [FWHM (full width at half maximum)]; ScR= scan rate for measurements in the IT; SR= scan range [m/z]; AGC= automatic gain control, max number of acquired ions per measurement; AcT= max. Ion acquisition time [ms]; CS= charge states used for fragmentation; IsM= Isolation mode (Q or IT), MS2 isolation and further is only done in IT; IsW= Isolation window [m/z], value followed by scan mode the isolation is based on (MS1, MS2 ...) Frag.= Fragmentation method; HCD= Higher-energy collisional dissociation; CID= Collision-induced dissociation; ETD= Electron-transfer dissociation; EThcD= Electron-Transfer/Higher-Energy Collision Dissociation; sHCD= stepped HCD; NCE= normalized collision energy; cycles: number of MSn recorded or max cycle time; RF= RF Lens [%]; SF= Source Fragmentation [V]; DDM: Data dependent Mode (cycle time in seconds, CT/[s] or number of scans, NS); NS= Number of data dependent scans.

**Table S6.**
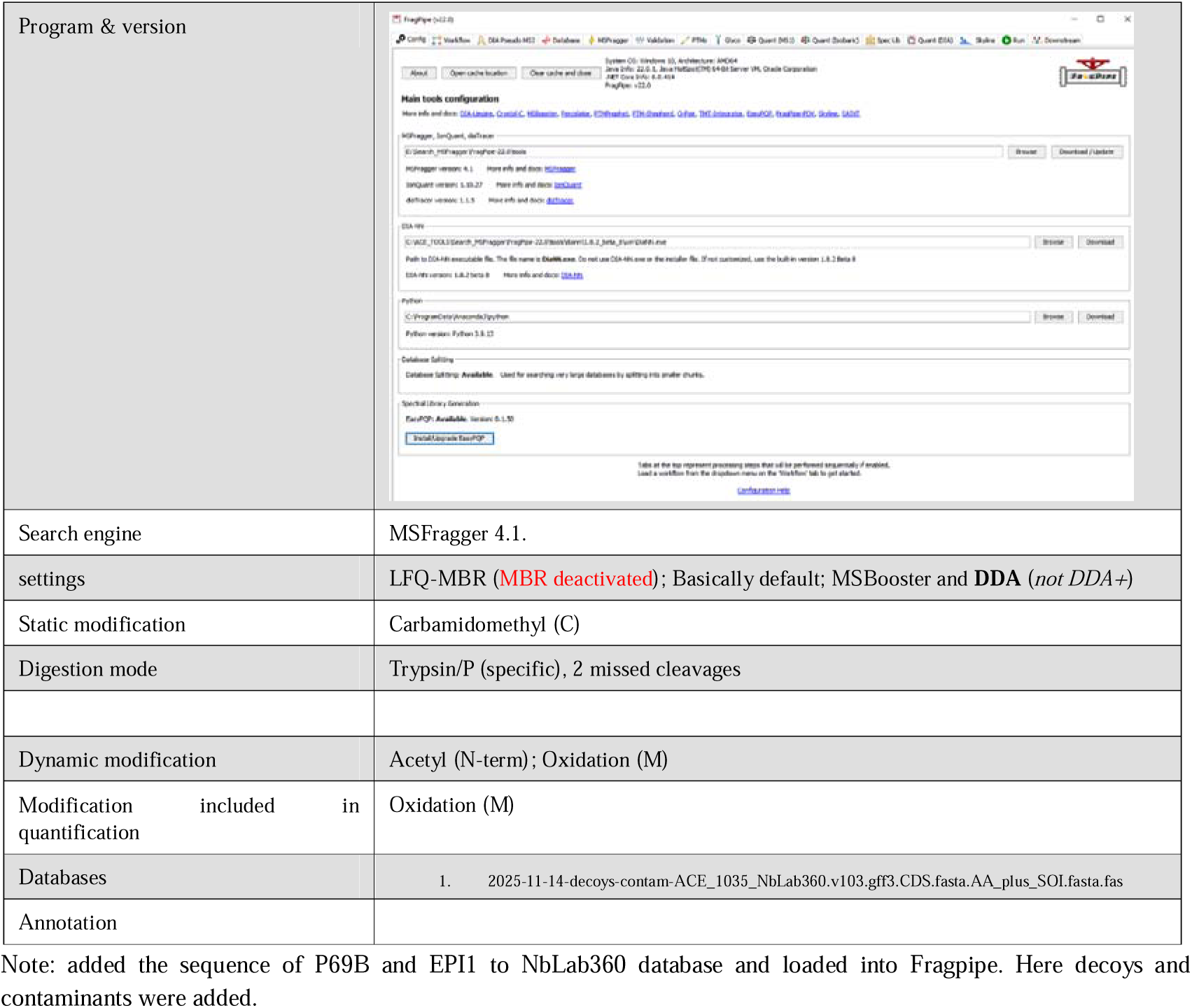
MSFragger search.

**Table S7.**
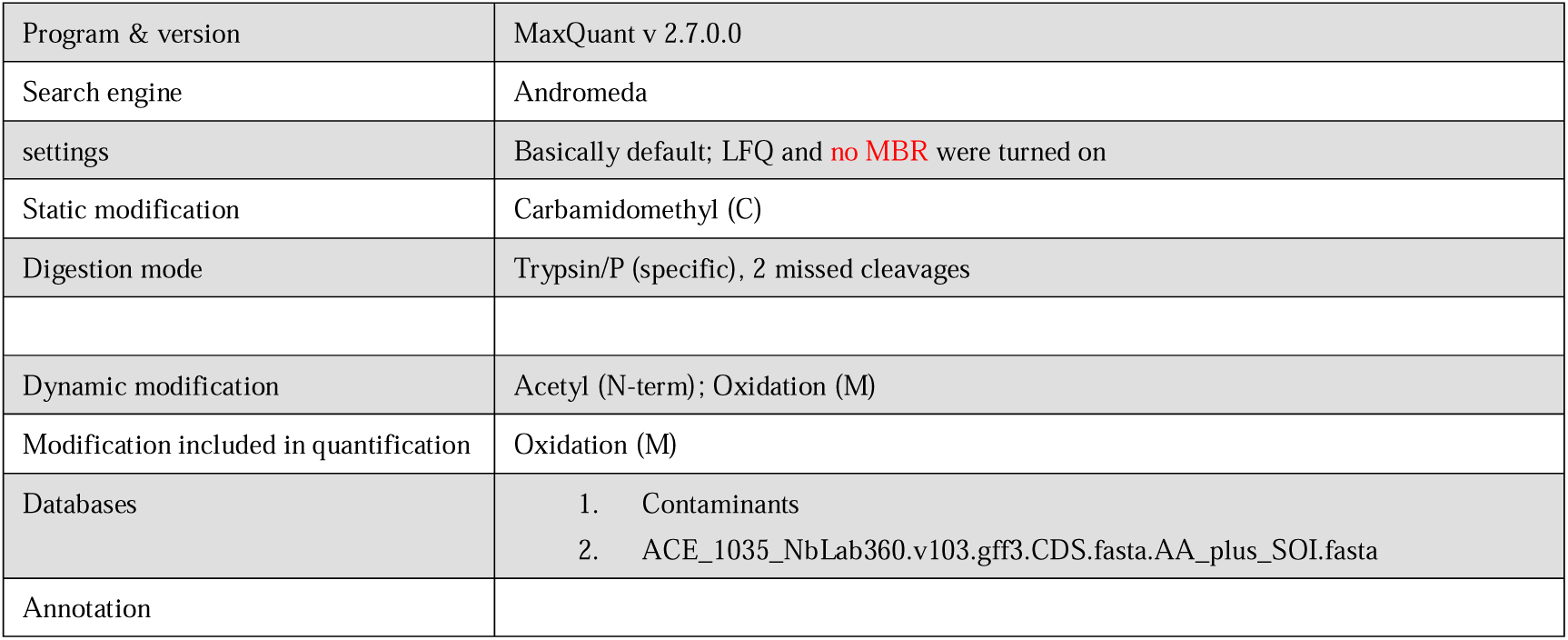
MaxQuant search.

## SUPPLEMENTAL FIGURES

**Fig. S1.**
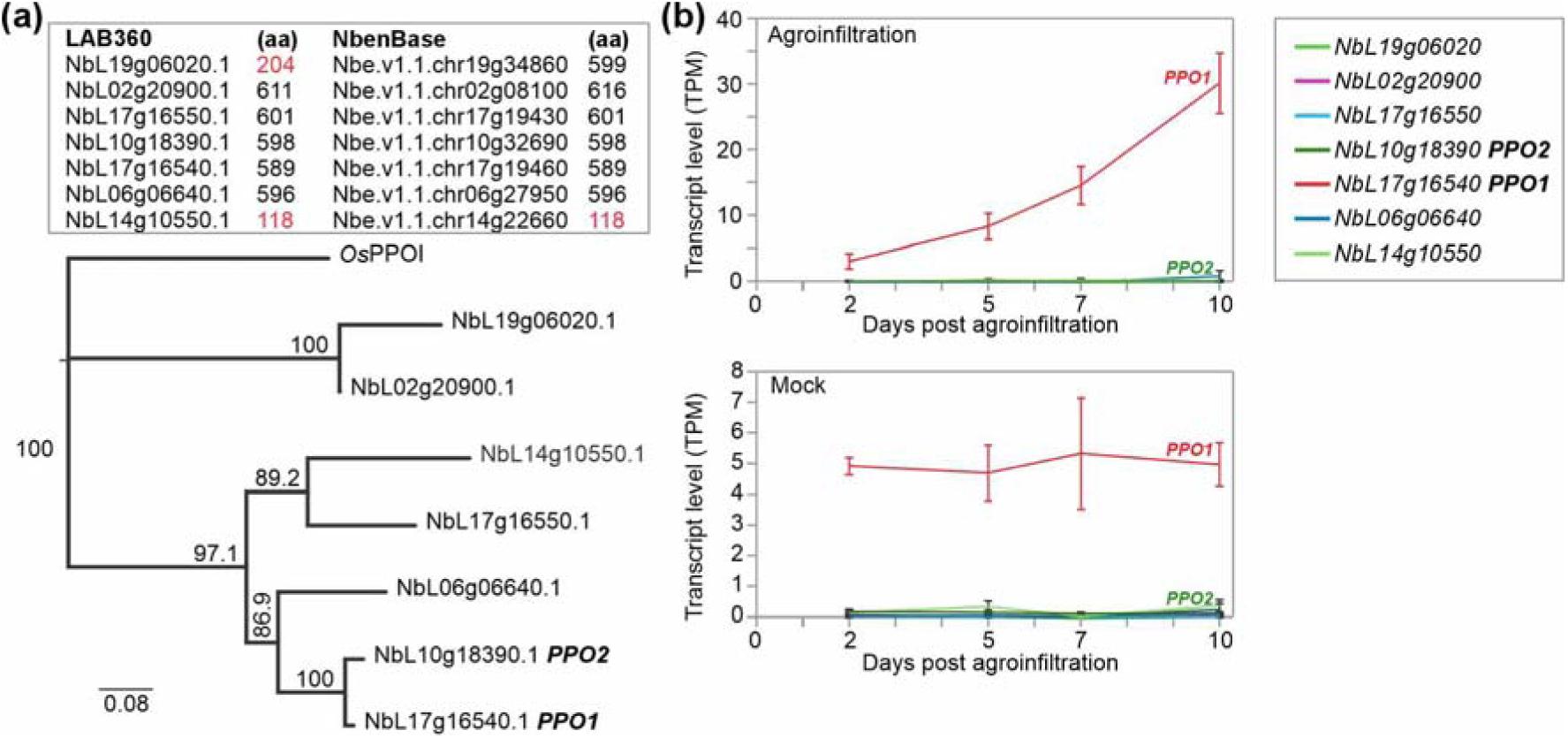
*PPO* genes in *Nicotiana benthamiana* **(a)** Upper panel: accession numbers of seven *PPO* genes from the LAB360 and NbenBase and the lengths of the proteins predicted from the open reading frames, showing a discrepancy only for the top listed gene. Lower panel: phylogeny of *PPO* genes. The tree was made with the Jukes–Cantor genetic distance model and the Neighbor-Joining method, with *O. sativa* PPOI as the outgroup. Branch support was assessed using 1000 bootstrap replicates. The analysis was performed in Geneious. **(b)** Transcript levels of the *PPO* genes in Mock and agroinfiltrated leaves in transcripts per million (TPM). Data were extracted from Grosse-Holz et al., 2018.

**Fig. S2.**
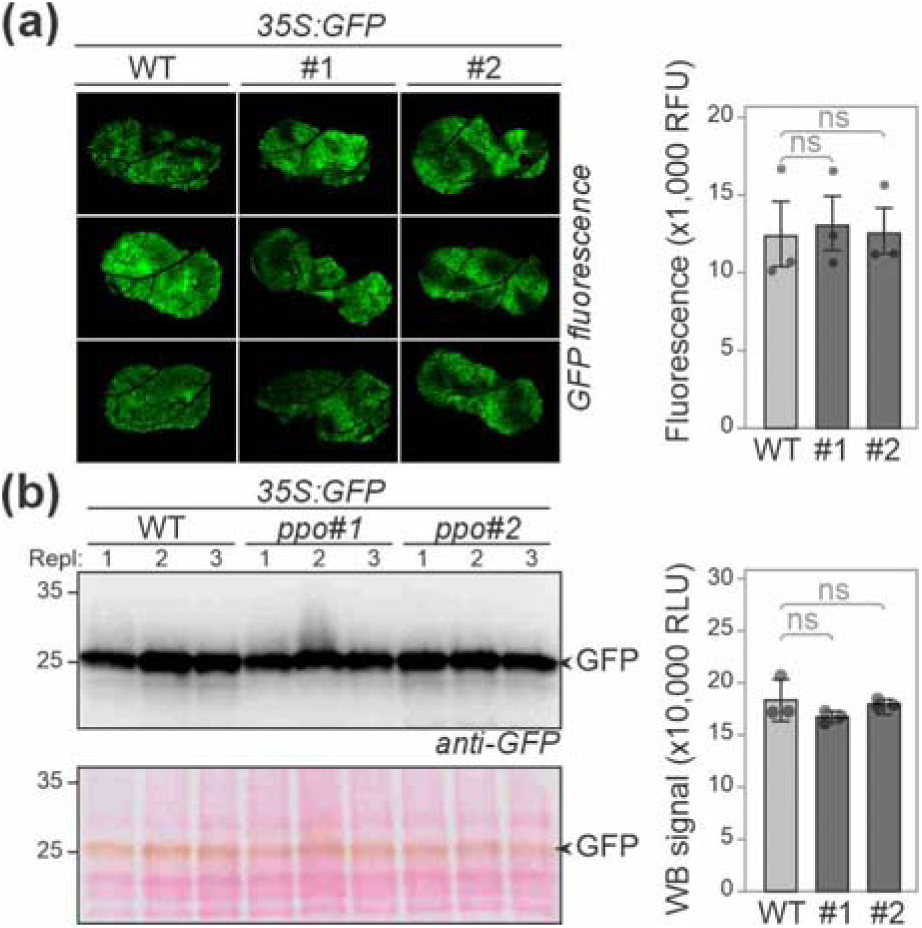
Transient protein expression is unaltered in *ppo* mutants. **(a)** Similar levels of GFP fluorescence upon transient expression in *ppo* mutant. Left: leaves were agroinfiltrated to transiently express *35S::GFP* and the fluorescence was imaged three days later (3dpi). Right: quantification of GFP fluorescence over three different plants. Error bars represent SE. P-values were calculated with ANOVA and found non-significant (ns, p>0.05). **(b)** Similar levels of GFP accumulation upon transient expression in *ppo* mutant. Left: leaf extracts generated from leaves transiently expressing GFP at 3dpi were analysed by anti-GFP western blot in n=3 biological replicates. Right: quantification of western signals. Error bars represent SE. P-values were calculated with ANOVA and found non-significant (ns, p>0.05).

**Fig. S3.**
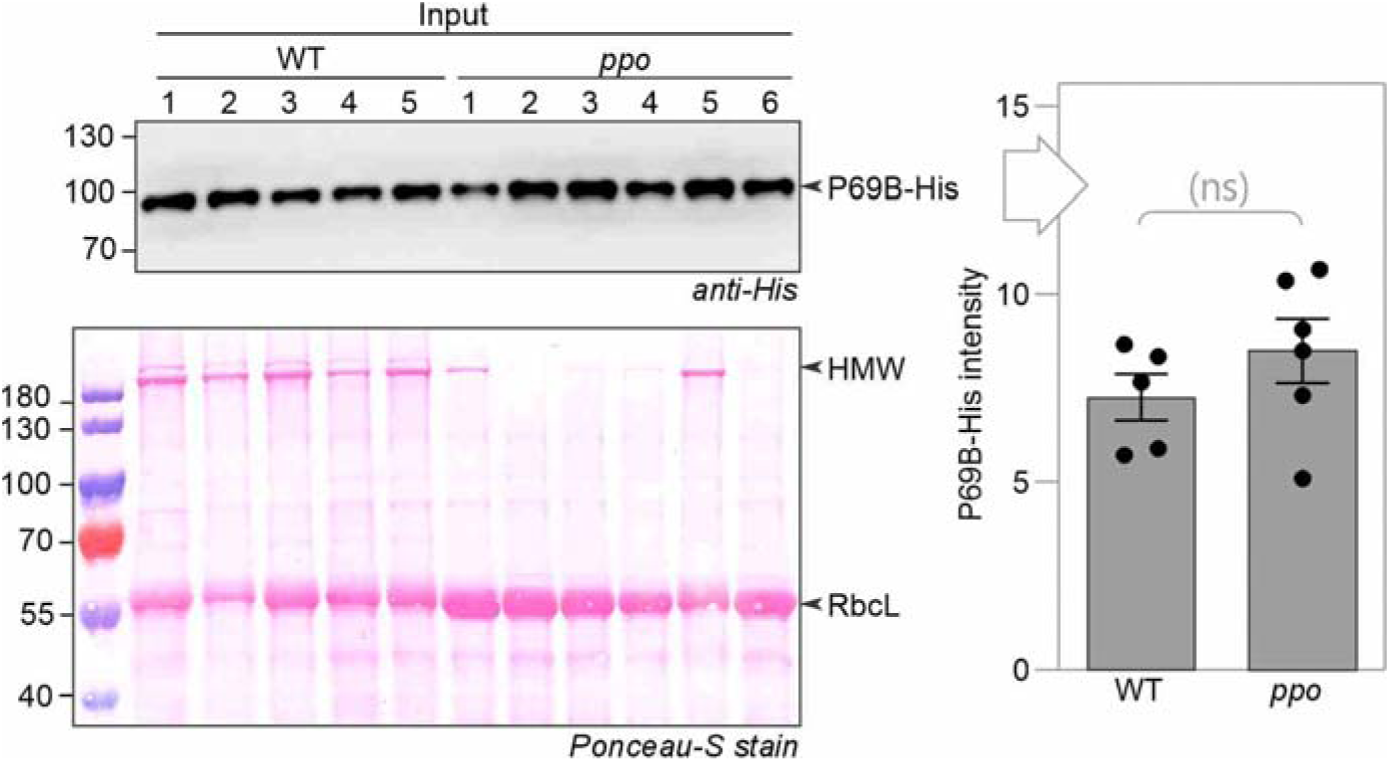
Input samples for P69B-His purification. Leaf total protein extracts of WT and *ppo* mutant line #1 transiently expressing P69B-His at 3dpi were analysed by western blot using the anti-P69B antibody and Ponceau-S staining. Western blot signals were quantified and plotted on the right. Error bars represent SE. P-values were determined with Student’s t-test and are non-significant (ns).

## References

Alamos S, Szarzanowicz MJ, Thompson MG, Stevens DM, Kirkpatrick LD, Dee A, Pannu H, Cui R, Liu S, Nimavat M, Krasileva K, Baidoo EEK, Shih PM. (2025) Quantitative dissection of Agrobacterium T-DNA expression in single plant cells reveals density-dependent synergy and antagonism. Nat Plants. 11: 1060–1073.

Bally J, Jung H, Mortimer C, Naim F, Philips JG, Hellens R, Bombarely A, Goodin MM, Waterhouse PM. (2018) The rise and rise of *Nicotiana benthamiana*: a plant for all reasons. Annual Review of Phytopathology 56: 405–426.

Beritza K, Watts EC, van der Hoorn RAL. (2024) Improving transient protein expression in agroinfiltrated Nicotiana benthamiana. New Phytol. 243: 846–850.

Beritza K, Buscaill P, Song S-J, Jutras PV, Huang J, Mach L, Dong S, van der Hoorn RAL. (2024) SBT5.2s are the major active extracellular subtilases processing IgG antibody 2F5 in the *Nicotiana benthamiana* apoplast. Plant Biotechnol J. 22: 2808–2810.

Bhonwong A, Stout MJ, Attajarusit J, Tantasawat P. (2009) Defensive role of tomato polyphenol oxidases against cotton bollworm (*Helicoverpa armigera*) and beet armyworm (*Spodoptera exigua*). J Chem Ecol. 35: 28–38.

Boeckx T, Winters AL, Webb KJ, Kingston-Smith AH. (2015) Polyphenol oxidase in leaves: is there any significance to the chloroplastic localization? J Exp Bot. 66: 3571–3579.

Busold S, Aglas L, Menage C, Auger L, Desgagnés R, Faye L, Fitchette A-C, de Jong EC, Martel C, Stigler M, Catala-Stordeur V, Tropper G, Vézina L-P, Gomord V, Geijtenbeek TBH, van Ree R. (2022) Fel d 1 surface expression on plant-made eBioparticles combines potent immune activation and hypoallergenicity. Allergy. 77: 3124–3126.

Busold S, Aglas L, Menage C, Auger L, Desgagnés R, Faye L, Fitchette A-C, de Jong EC, Martel C, Stigler M, Catala-Stordeur V, Tropper G, Vézina L-P, Gomord V, Geijtenbeek TBH, van Ree R. (2024) Plant-produced *Der p* 2-bearing bioparticles activate Th1/Treg-related activation patterns in dendritic cells irrespective of the allergic background. Clin Exp Allergy. 54: 300–303.

Buyel JF. (2025) Developing downstream processes for the purification of recombinant proteins and small molecules from *Nicotiana benthamiana* biomass. Plant Biotechnol J. in press

Charland N, Gobeil P, Pillet S, Boulay I, Séguin A, Makarkov A, Heizer G, Bhutada K, Mahmood A, Trépanier S, Hager KJ, Jiang-Wright J, Atkins J, Saxena P, Cheng MP, Vinh DC, Boutet P, Roman F, Van Der Most R, Ceregido MA, Dionne M, Tellier G, Gauthier J-S, Essink B, Libman M, Haffizulla J, Fréchette A, D’Aoust M-A, Landry N, Ward BJ. (2022) Safety and immunogenicity of an AS03-adjuvanted plant-based SARS-CoV-2 vaccine in Adults with and without Comorbidities. npj Vaccines. 7:142.

Constabel CP, Bergey DR, Ryan CA. (1995) Systemin activates synthesis of wound-inducible tomato leaf polyphenol oxidase via the octadecanoid defense signaling pathway. Proc Natl Acad Sci U S A. 92: 407–411.

Daduang R, Suwanchaikasem P, Rattanapisit K, Vitayathikornnasak S, Srisangsung T, Bulaon CJI, Phoolcharoen W. (2025) LC-MS determination of *Nicotiana benthamiana* host plant proteins in the drug products of recombinant plant-produced pembrolizumab. Sci Rep. 15: 25635

de Taeye SW, Faye L, Morel B, Schriek AI, Umotoy JC, Yuan M, Kuzmina NA, Turner HL, Zhu X, Grünwald-Gruber C, Poniman M, Burger JA, Caniels TG, Fitchette AC, Desgagnés R, Stordeur V, Mirande L, Beauverger G, de Bree G, Ozorowski G, Ward AB, Wilson IA, Bukreyev A, Sanders RW, Vezina LP, Beaumont T, van Gils MJ, Gomord V. (2025) Plant-produced SARS-CoV-2 antibody engineered towards enhanced potency and in *vivo* efficacy. Plant Biotechnol J. 23: 4–16.

Demichev V, Messner CB, Vernardis SI, Lilley KS, Ralser M. (2019) DIA-NN: neural networks and interference correction enable deep proteome coverage in high throughput. Nat Methods. 17: 41–44.

Diao HP, Meng HX, Xu XJ, Zhang ZL, Zhang JF, Guo YF, Hwang I, Song SJ. (2026) Knocking out two polyphenol oxidase genes significantly improves recombinant protein purification in *Nicotiana benthamiana*. Plant Biotechnol J. 10.1111/pbi.70575

Dodds IL, Watts EC, Schuster M, Buscaill P, Tumas Y, Holton NJ, Song S, Stuttmann J, Joosten MHAJ, Bozkurt T, van der Hoorn RAL. (2025) Immunity gene silencing increases transient protein expression in Nicotiana benthamiana. Plant Biotechnol J. 23: 1768–1770.

Dudley QM, Jo S, Guerrero DAS, Chhetry M, Smedley MA, Harwood WA, Sherden NH, O’Connor SE, Caputi L, Patron NJ. (2022) Reconstitution of monoterpene indole alkaloid biosynthesis in genome-engineered *Nicotiana benthamiana*. Commun Biol. 5: 949.

Felton GW, Donato K, Del Vecchio RJ, Duffey SS. (1989) Activation of plant foliar oxidases by insect feeding reduces nutritive quality of foliage for noctuid herbivores. J Chem Ecol. 15:2667–2694.

Golubova D, Tansley C, Su H, Patron NJ. (2024) Engineering *Nicotiana benthamiana* as a platform for natural product biosynthesis. Curr Opin Plant Biol. 81: 102611.

González MN, Massa GA, Andersson M, Turesson H, Olsson N, Fält A-S, Storani L, Décima Oneto CA, Hofvander P, Feingold SE. (2020) Reduced enzymatic browning in potato tubers by specific editing of a polyphenol oxidase gene via ribonucleoprotein complexes delivery of the CRISPR/Cas9 system. Front Plant Sci. 10: 1649.

Grosse-Holz F, Kelly S, Blaskowski S, Kaschani F, Kaiser M, van der Hoorn RAL. (2018) The transcriptome, extracellular proteome and active secretome of agroinfiltrated *Nicotiana benthamiana* uncover a large, diverse protease repertoire. Plant Biotechnol J. 16: 1068–1084.

Homma F, Huang J, van der Hoorn RAL. (2023) AlphaFold-Multimer predicts cross-kingdom interactions at the plant-pathogen interface. Nat Commun. 14: 6040.

Hager KJ, Pérez Marc G, Gobeil P, Diaz RS, Heizer G, Llapur C, Makarkov AI, Vasconcellos E, Pillet S, Riera F, Saxena P, Geller Wolff P, Bhutada K, Wallace G, Aazami H, Jones CE, Polack FP, Ferrara L, Atkins J, Boulay I, Dhaliwall J, Charland N, Couture MMJ, Jiang-Wright J, Landry N, Lapointe S, Lorin A, Mahmood A, Moulton LH, Pahmer E, Parent J, Séguin A, Tran L, Breuer T, Ceregido MA, Koutsoukos M, Roman F, Namba J, D’Aoust MA, Trepanier S, Kimura Y, Ward BJ. (2022) CoVLP Study Team. Efficacy and safety of a recombinant plant-based adjuvanted Covid-19 vaccine. N Engl J Med. 386: 2084–2096.

Hughes CS, Moggridge S, Müller T, Sorensen PH, Morin GB, Krijgsveld J. (2019) Single-pot, solid-phase-enhanced sample preparation for proteomics experiments. Nat Protoc. 14: 68–85.

Jutras PV, Dodds I, van der Hoorn RAI. (2020) Proteases of *Nicotiana benthamiana*: an emerging battle for molecular farming. Curr Opin Biotechnol. 61: 60–65.

Kaldis A, Uddin MS, Guluarte JO, Martin C, Alexander TW, Menassa R. (2023) Development of a plant-based oral vaccine candidate against the bovine respiratory pathogen *Mannheimia haemolytica*. Frontiers in Plant Science. 14: 1251046.

Kaschani F, Gu C, Niessen S, Hoover H, Cravatt BF, van der Hoorn RAL. (2009) Diversity of serine hydrolase activities of unchallenged and botrytis-infected *Arabidopsis thaliana*. Mol. Cell. Proteomics 8: 1082–1093.

Kong AT, Leprevost FV, Avtonomov DM, Mellacheruvu D, Nesvizhskii AI. (2017) MSFragger: ultrafast and comprehensive peptide identification in mass spectrometry-based proteomics. Nat Methods. 14: 513–520.

Kurotani KI, Hirakawa H, Shirasawa K, Tanizawa Y, Nakamura Y, Isobe S, Notaguchi M. (2023) Genome sequence and analysis of *Nicotiana benthamiana*, the model plant for interactions between organisms. Plant Cell Physiol. 64: 248–257.

Lawson AW, Macha A, Neumann U, Gunkel M, Chai J, Behrmann E, Schulze-Lefert P. (2025) Purifying recombinant proteins from *Nicotiana benthamiana* for structural studies. Nat Protoc. in press

Mahadevan C, Watts EC, Zheng K, Song SJ, van der Hoorn RAL. (2025) Polyphenol oxidase silencing avoids protein cross-linking and enzymatic browning in *Nicotiana benthamiana* leaf extracts. Plant Biotechnol J. 24: 96–98.

Ranawaka B, An J, Lorenc MT, Jung H, Sulli M, Aprea G, Roden S, Llaca V, Hayashi S, Asadyar L, LeBlanc Z, Ahmed Z, Naim F, de Campos SB, Cooper T, de Felippes FF, Dong P, Zhong S, Garcia-Carpintero V, Orzaez D, Dudley KJ, Bombarely A, Bally J, Winefield C, Giuliano G, Waterhouse PM. (2023) A multi-omic *Nicotiana benthamiana* resource for fundamental research and biotechnology. Nat Plants. 9: 1558–1571.

Rappsilber J, Mann M, Ishihama Y. (2007) Protocol for micro-purification, enrichment, pre-fractionation and storage of peptides for proteomics using StageTips. Nat Protoc. 2: 1896–906.

Ren Z, Liu Y, Huang J, An L, Zhang Y, Yang W, Lei T. (2024) Low oxygen concentration alleviates banana peel browning by inhibiting membrane lipid oxidation and polyphenol oxidase activity. Int J Food Sci Technol. 59: 3350–3359.

Schuster M, Paulus JK, Kourelis J, van der Hoorn RAL. (2022) Purification of His-tagged proteases from the apoplast of agroinfiltrated *N. benthamiana*. Methods Mol Biol. 2447: 53–66.

Shanmugaraj B, Rattanapisit K, Manopwisedjaroen S, Thitithanyanont A, Phoolcharoen W. (2020) Monoclonal antibodies B38 and H4 produced in *Nicotiana benthamiana* neutralize SARS-CoV-2 in vitro. Front Plant Sci. 11: 589995.

Sommer A, Ne’eman E, Steffens JC, Mayer AM, Harel E. (1994) Import, targeting, and processing of a plant polyphenol oxidase. Plant Physiol. 105: 1301–1311.

Sui X, Meng Z, Dong T, Fan X, Wang Q. (2023) Enzymatic browning and polyphenol oxidase control strategies. Curr. Opin. Biotechnol. 81: 102921.

Tang M-G, Zhang S, Xiong L-G, Zhou J-H, Huang J-A, Zhao A-Q, Liu Z-H, Liu A-L. (2023) A comprehensive review of polyphenol oxidase in tea (*Camellia sinensis*): Physiological characteristics, oxidation manufacturing, and biosynthesis of functional constituents. Compr Rev Food Sci Food Saf. 22: 2267–2291.

Terefe NS, Buckow R, Versteeg C. (2014) Quality-related enzymes in fruit and vegetable products: effects of novel food processing technologies, part 1: high-pressure processing. Crit Rev Food Sci Nutr. 54: 24–63.

Thipyapong P, Hunt MD, Steffens JC. (2004) Antisense downregulation of polyphenol oxidase results in enhanced disease susceptibility. Planta. 220: 105–117.

Tyanova S, Temu T, Sinitcyn P, Carlson A, Hein MY, Geiger T, Mann M, Cox J. (2016) The Perseus computational platform for comprehensive analysis of (prote)omics data. Nat Methods. 13: 731–40.

VanderBurgt JT, Harper O, Garnham CP, Kohalmi SE, Menassa R. (2023) Plant production of a virus-like particle-based vaccine candidate against porcine reproductive and respiratory syndrome. Front. Plant Sci. 14: 1044675.

Vizcaíno JA, Csordas A, del-Toro N, Dianes JA, Griss J, Lavidas I, Mayer G, Perez-Riverol Y, Reisinger F, Ternent T, Xu QW, Wang R, Hermjakob H. (2016) 2016 update of the PRIDE database and its related tools. Nucleic Acids Res. 44: D447–56.

Vollheyde K, Dudley QM, Yang T, Oz MT, Mancinotti D, Fedi MO, Heavens D, Linsmith G, Chhetry M, Smedley MA, Harwood WA, Swarbreck D, Geu-Flores F, Patron NJ. (2023) An improved *Nicotiana benthamiana* bioproduction chassis provides novel insights into nicotine biosynthesis. New Phytol. 240: 302–317.

Ward BJ, Makarkov A, Séguin A, Pillet S, Trépanier S, Dhaliwall J, Libman MD, Vesikari T, Landry N. (2020) Efficacy, immunogenicity, and safety of a plant-derived, quadrivalent, virus-like particle influenza vaccine in adults (18–64 years) and older adults (≥65 years): two multicentre, randomised phase 3 trials. Lancet. 396: 1491–1503.

Ward BJ, Gobeil P, Séguin A, Atkins J, Boulay I, Charbonneau P-Y, Couture M, D’Aoust M-A, Dhaliwall J, Finkle C, Hager KJ, Mahmood A, Makarkov A, Cheng MP, Pillet S, Schimke PA, St-Martin S, Trépanier S, Landry N. (2021) Phase 1 randomized trial of a plant-derived virus-like particle vaccine for COVID-19. Nat Med. 27: 1071–1078.

Win J, Kamoun S, Jones AM. (2011) Purification of effector-target protein complexes via transient expression in *Nicotiana benthamiana*. Methods Mol Biol. 712: 181–94.

Yang KL, Yu F, Teo GC, Li K, Demichev V, Ralser M, Nesvizhskii AI. (2023) MSBooster: improving peptide identification rates using deep learning-based features. Nat Commun. 14: 4539.

Zhang J, Sun X. (2021) Recent advances in polyphenol oxidase-mediated plant stress responses. Phytochemistry. 179: 112588.

Zhang S. (2023) Recent Advances of Polyphenol Oxidases in Plants. Molecules. 28: 2158.

Zheng K, Lyu J, Thomas EL, Schuster M, Sanguankiattichai N, Ninck S, Kaschani F, Kaiser M, van der Hoorn RAL. (2024) The proteome of *Nicotiana benthamiana* is shaped by extensive protein processing. New Phytol. 243: 1034–1049.

Zou H, Li C, Wei X, Xiao Q, Tian X, Zhu L, Ma B, Ma F, Li M. (2025) Expression of the polyphenol oxidase gene *MdPPO7* is modulated by *MdWRKY3* to regulate browning in sliced apple fruit. Plant Physiol. 197: kiae614.

## Supplemental references

Paulus JK, Kourelis J, Ramasubramanian S, Homma F, Godson A, Hörger AC, Hong TN, Krahn D, Ossorio Carballo L, Wang S, Win J, Smoker M, Kamoun S, Dong S, van der Hoorn RAL. (2020) Extracellular proteolytic cascade in tomato activates immune protease Rcr3. Proc Natl Acad Sci USA. 117: 17409–17417.

Van Der Hoorn RAL, Rivas S, Wulff BB, Jones JDG, Joosten MHAJ. (2003) Rapid migration in gel filtration of the Cf-4 and Cf-9 resistance proteins is an intrinsic property of Cf proteins and not because of their association with high-molecular-weight proteins. Plant J. 35: 305–315.

